# Sequence-encoded differences in the conformational ensembles of CITED transcriptional activation domains impact coactivator binding

**DOI:** 10.64898/2026.01.20.700670

**Authors:** To Uyen Do, Emma J. Kraft, Garrett F. Chappell, Stuart Parnham, Rebecca B. Berlow

## Abstract

Recent advances in predicting and modeling conformational ensembles of intrinsically disordered proteins (IDPs) have provided much needed insights into sequence-ensemble relationships. It is thought that conservation of physicochemical properties, but not the exact identity or order of the amino acids, maintains IDP ensemble properties that are crucial for function. However, detailed experimental studies are still required to fully understand the relationships between sequence and function in IDPs. The human CITED proteins, which are fully disordered transcriptional regulators, share conserved C-terminal transactivation domains (CTADs) that interact with the TAZ1 domain of the transcriptional coactivators CBP/p300. The conserved CTADs harbor amino acid substitutions in regions that are known to be important for interactions of CITED2 with TAZ1, but the effects of these substitutions on TAZ1 binding for the other CITED proteins are unknown. Here, we use solution NMR spectroscopy, circular dichroism, and surface plasmon resonance to characterize the conformational ensembles, dynamics, and interactions of the CITED CTADs. The CTADs are disordered in isolation, although the CITED2 CTAD uniquely displays residual helical structure that is sensitive to ionic strength and protein concentration. In contrast, the CITED1 and CITED4 CTADs remain largely disordered and exhibit more uniform dynamics. Quantitative binding measurements reveal differences in thermodynamics and kinetics for the CTADs’ interactions with TAZ1, with CITED2 binding most tightly and CITED4 exhibiting significantly weaker affinity. Our results highlight the sensitivity of IDP conformational ensembles to minor sequence changes and the impacts that changes in IDP structures and dynamics can have on biological functions.

## Introduction

Intrinsically disordered proteins (IDPs) have important roles in nearly every facet of biology^1,2^. The intrinsic flexibility and lack of stable structure in IDPs confers unique functional advantages, particularly in critical regulatory processes that must be both robust and efficient^3^. Research over the past few decades has provided insights into the numerous roles of IDPs in transcriptional regulation, with a particular focus on mapping interactions of a wide array of transcription factors and transcriptional regulators that use intrinsically disordered regions (IDRs) to interact with other components of the transcriptional machinery.

Two of the most studied systems for decoding disordered protein interactions in transcriptional regulation are the general transcriptional coactivators CREB Binding Protein (CBP) and its paralog p300^4,5^. CBP and p300 are partially disordered but their folded domains are implicated as binding hubs for numerous disordered partners^1,4,6,7^. Interactions between CBP/p300 and other molecular partners are typically mediated by intrinsically disordered regions known as transcriptional activation domains^8^. Transcriptional activation domains are generally found in transcription factors, where they function in concert with DNA binding domains to bind to cognate sites on DNA, but can also be found in proteins that lack DNA binding activity but have important functions in regulating transcriptional processes.

The CITED (<u>**C**</u>BP/p300-<u>**I**</u>nteracting <u>**T**</u>ransactivator with <u>**ED**</u>-rich tail) family of proteins (CITED1, 2, and 4 in humans) are fully disordered transcriptional regulators with partially overlapping biological functions^9–15^. The CITED proteins share a C-terminal transactivation domain (CTAD) of ~50 residues that, as their name implies, is enriched in glutamate and aspartate residues. Previous work has described the interactions of the CITED2 CTAD with the TAZ1 domain of CBP^16–20^. The CITED2 CTAD binds to TAZ1 in a multivalent manner, with three highly conserved motifs (the *α*A helix, LPEL, and ϕC motifs) that function synergistically within the polypeptide chain to bind to TAZ1 with a *K*_d_ of 10 nM^18–20^. The CITED proteins have all been reported to bind CBP/p300, but the exact mechanisms and sequence determinants for these interactions are not known^13,21,22^.

While IDP sequences are generally variable and challenging to interpret using traditional bioinformatics methods^2^, the CITED CTAD sequences retain a high degree of similarity overall (Figure 1). Yet, the CTAD sequences of CITED1 and CITED4 harbor amino acid substitutions in regions that are known to be important for CITED2 CTAD:TAZ1 interactions. As the conformational ensembles of IDPs are dictated by amino acid sequence, we hypothesized that variations in the sequences could be important for encoding functional differences in the CITED CTADs.

**Figure 1.**
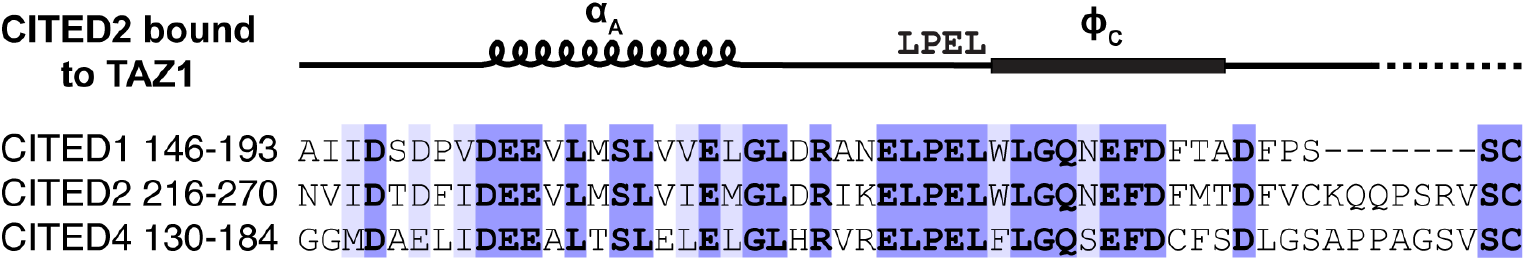
Sequence alignment of the C-terminal transactivation domains of human CITED1, 2, and 4. Identical residues among the three sequences are in bold font and highlighted in blue. Positions with conservative substitutions are highlighted in light blue. CITED2 binding motifs that are important for TAZ1 interactions are annotated above the sequence.

Here, we use a combination of biophysical approaches (including solution NMR spectroscopy, circular dichroism (CD) spectroscopy, and surface plasmon resonance) to characterize the conformational ensembles, dynamics, and interactions of the CITED CTADs. We find that differences in the sequences of these domains encode differences in structural propensities, overall dynamics, and interactions with TAZ1. Our results highlight the importance of the exquisite sensitivity of IDP conformational ensembles to minor sequence changes and the impacts that alteration of IDP structures and dynamics can have on requisite biological functions.

## Results

We generated sequence alignments of the CITED CTAD sequences to identify similarities and differences between the human CITED CTAD sequences (Figure 1) and across species (Supplementary Figures 1, 2, and 3). Comparison of the CITED CTAD sequences reveals that they are highly conserved, particularly in comparison to other disordered protein sequences. The sequences of the CITED1 and CITED4 CTADs are 45% identical to the CITED2 CTAD. Outside of these identical regions, there are an additional 7 positions (13%) that feature conservative substitutions in CITED1 and/or CITED4. The identical residues of CITED1 and CITED4 align with positions in the CITED2 CTAD that are known to be important for high-affinity binding to the TAZ1 domain of the general transcriptional coactivator CBP^16–19^. For instance, 60% of the residues in the CITED2 *α*A helix (residues 225-235) are identical in CITED1 and CITED4, while an even higher degree of conservation is observed for the LPEL motif (CITED2 residues 243-246) and for residues C-terminal to the LPEL motif that make stabilizing contacts in the CITED2:TAZ1 complex (Figure 1). These trends in conservation are even more pronounced when comparing sequences of the individual CTADs across species (Supplementary Figures 1, 2, and 3), suggesting that the sequence differences in the CITED CTADs have been preserved across evolution.

**Figure 2.**
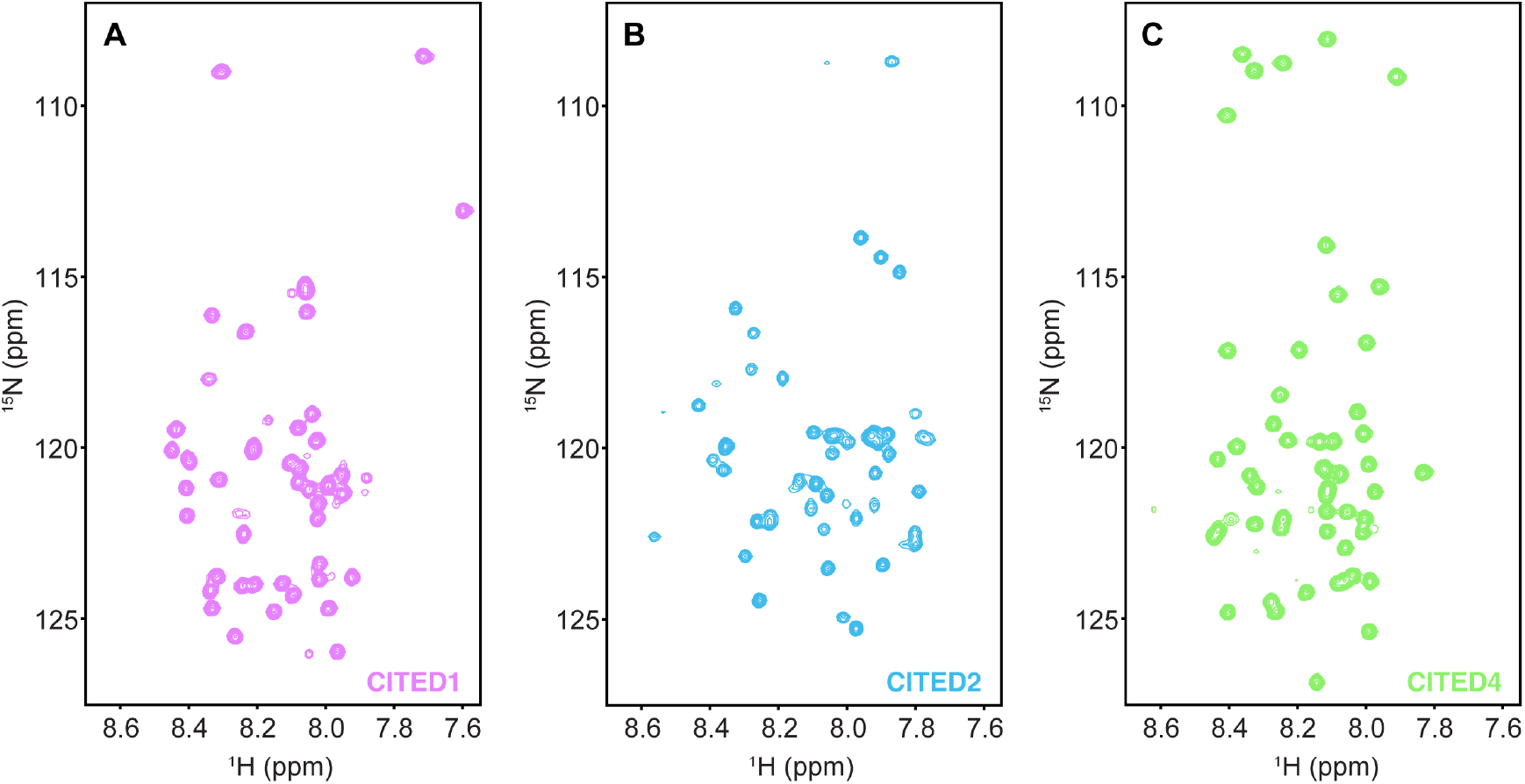
The CITED CTADs are intrinsically disordered. ^1^H-^15^N HSQC spectra are shown for the ^15^N-labeled CITED1 (A), CITED2 (B), and CITED4 (C) CTADs. All spectra were collected for 100 µM samples at 25 °C in buffer containing 20 mM Tris-HCl pH 6.8, 150 mM NaCl, 2 mM DTT, and 1% D_2_O. Spectra are all plotted at the same contour level for ease of comparison.

**Figure 3.**
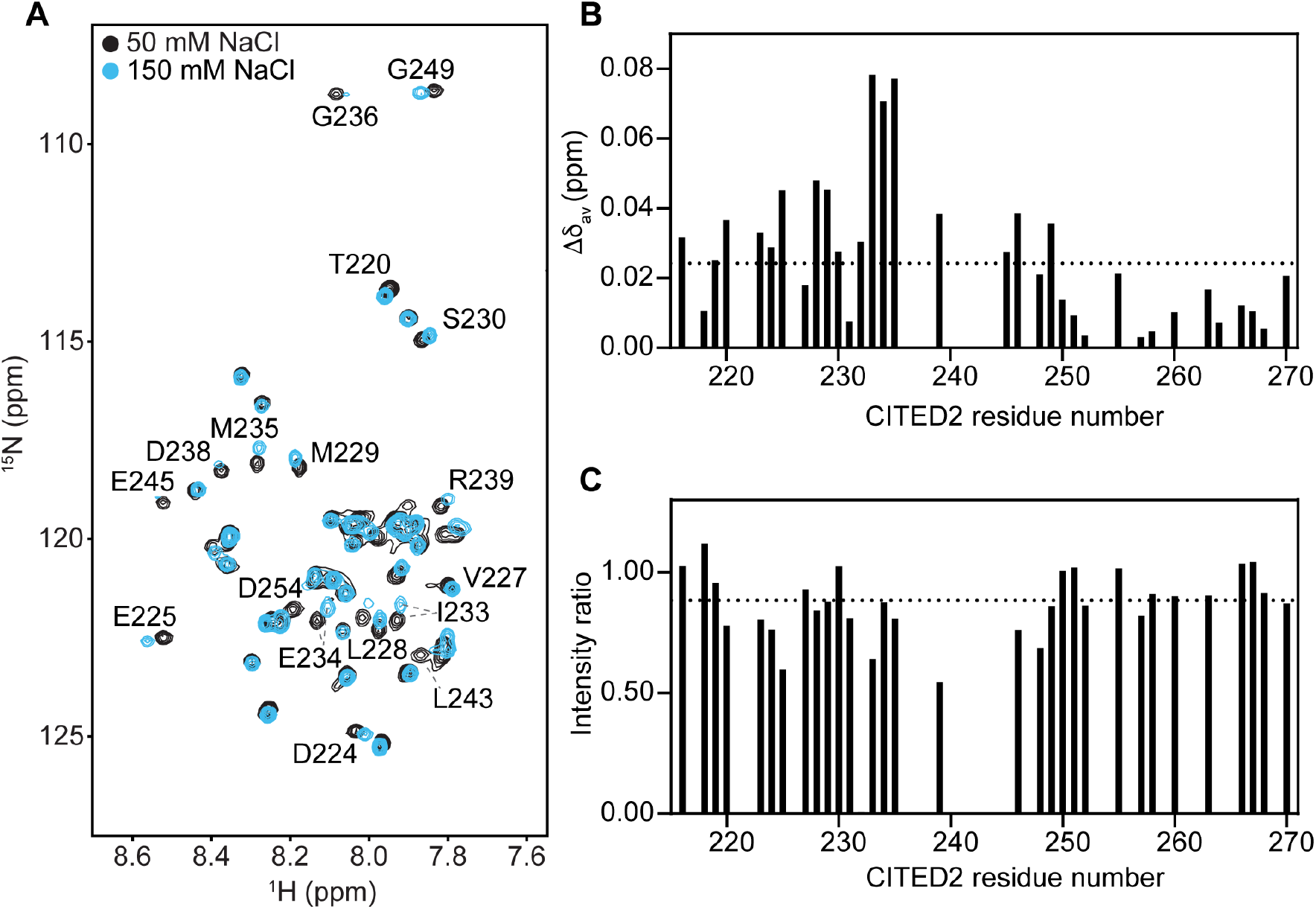
The CITED2 CTAD shows ionic strength-dependent peak shifts and broadening. (A) Superimposed ^1^H-^15^N HSQC spectra of 100 µM ^15^N-CITED2 CTAD in buffer containing 50 mM NaCl (black) or 150 mM NaCl (light blue) and 20 mM Tris-HCl pH 6.8, 2 mM DTT, and 1% D_2_O. Both spectra were collected at 25 °C. (B) Weighted average ^1^H-^15^N chemical shift changes (Δδav) for each residue in the CITED2 CTAD for spectra collected in buffer containing 50 mM and 150 mM NaCl. (C) Ratio of peak intensities for CITED2 CTAD residues from spectra collected at different NaCl concentrations. Intensity ratios were calculated by dividing the peak intensities from the 150 mM NaCl spectrum by the peak intensities from the 50 mM NaCl spectrum. In (B) and (C), data are omitted for resonances that are broadened beyond detection or overlapped in at least one of the spectra. Dashed horizontal lines represent the 10% trimmed mean.

We hypothesized that these evolutionarily conserved differences in the sequences of the CITED CTADs could have impacts on their conformational ensembles. The CITED CTADs are all predicted to be intrinsically disordered, and disorder of the CITED2 CTAD was previously confirmed using solution NMR spectroscopy^17^. To the best of our knowledge, the CITED1 and CITED4 CTADs have never been studied by biophysical methods, and data for the CITED2 CTAD in its unbound state is limited.

To further investigate the structural properties of the CITED CTADs, we expressed and purified ^15^N-labeled proteins for NMR studies. ^1^H-^15^N HSQC spectra confirm that the CITED CTADs are all disordered (Figure 2), with the ^1^H chemical shifts of all observable resonances falling into a relatively narrow chemical shift range centered around ~8.1 ppm. This limited ^1^H chemical shift dispersion for amide backbone resonances is characteristic of disordered proteins. However, further inspection of the spectra highlights subtle differences in the CITED2 spectrum in comparison to the spectra of CITED1 and CITED4. For CITED1 and CITED4, the peaks are well-resolved and relatively uniform in intensity, suggesting that these proteins are highly dynamic and unstructured (Figure 2, A and C). In contrast, the peaks in the CITED2 spectrum are variable in intensity and occupy a slightly wider range of ^1^H chemical shifts, suggesting that there may be some residual structure in the CITED2 CTAD that is partially restricting the dynamics of selected residues (Figure 2B).

Potential differences in dynamics for the CITED2 CTAD are also supported by the linewidths of the amide resonances. Overall, the ^15^N linewidths of CITED2 resonances are higher and more variable than the linewidths in the CITED1 and CITED4 spectra. For the CITED2 spectra, the mean and standard deviation of the ^15^N linewidths for all resonances is 23 ± 3 Hz. The CITED1 and CITED4 resonances have narrower ^15^N linewidths (18 ± 1 Hz for both spectra). These trends are consistent with the observable intensity differences in the spectra and highlight that there are unique structural and dynamic features in the CITED2 CTAD in comparison with CITED1 and CITED4.

We assigned the backbone chemical shifts of the CITED2 CTAD so that we could identify which residues of CITED2 are required for the observed structural and dynamic changes. Using standard triple-resonance methods, we were able to assign 98% of the ^1^H, ^13^C, and ^15^N backbone chemical shifts for the CITED2 CTAD (Supplementary Figure 4). As we were optimizing sample conditions for completing the backbone chemical shift assignments, we observed that the CITED2 CTAD spectrum is sensitive to changes in buffer conditions, including changes in ionic strength and protein concentration. Changing the salt concentration of the buffer has non-uniform effects on the CITED2 CTAD resonances (Figure 3). Many of the larger differences in chemical shift and intensity for spectra collected at 50 mM NaCl and 150 mM NaCl are observed for resonances that correspond to CITED2 residues in the *α*A helix. There are also significant chemical shift perturbations for hydrophobic and polar residues in the ϕC region. In contrast, there are much smaller changes in the spectra of the CITED1 and CITED4 CTADs in response to changing salt concentration (Supplementary Figure 5), highlighting that this behavior is a unique feature of the CITED2 CTAD.

**Figure 4.**
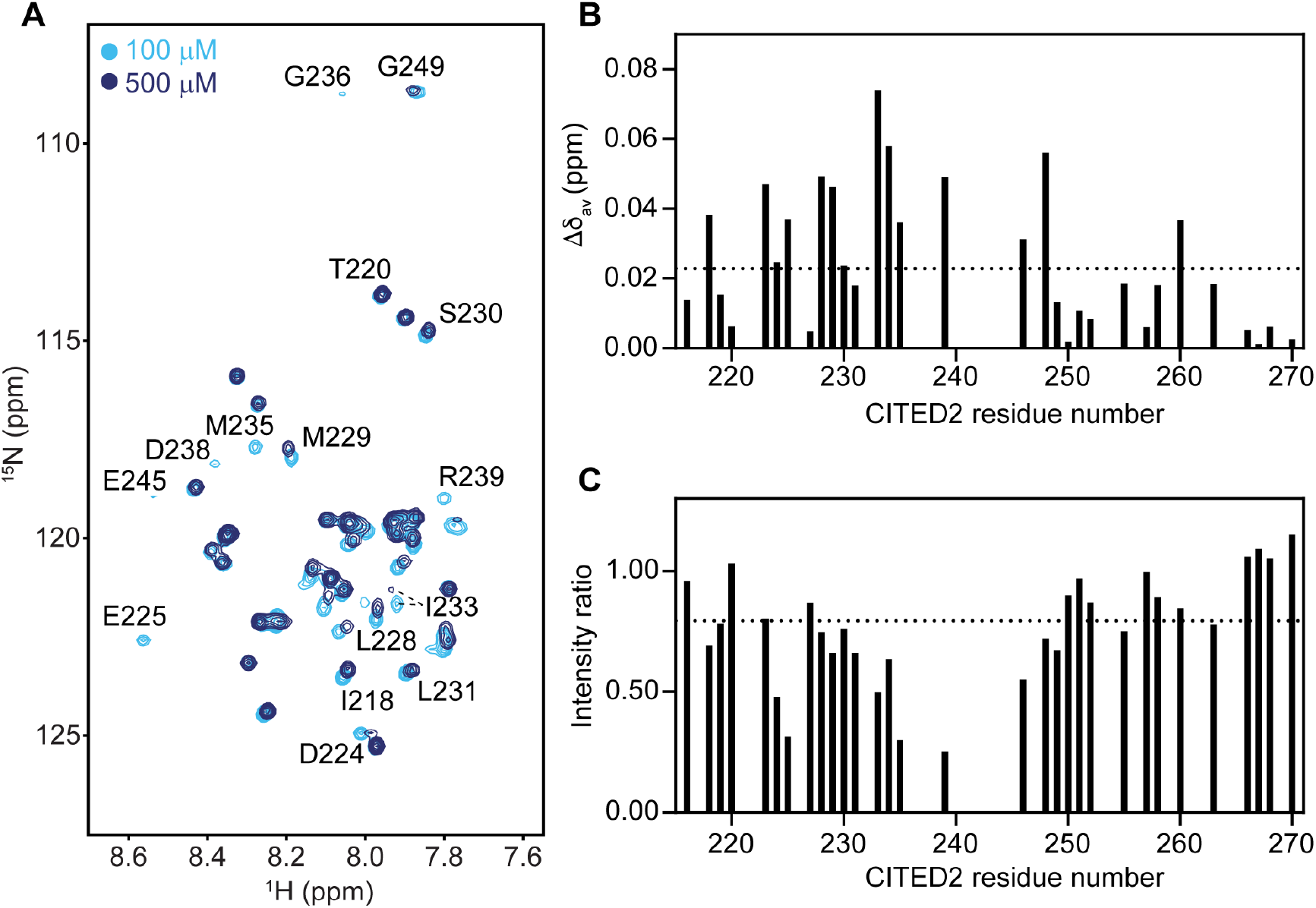
NMR spectra of the CITED2 CTAD are concentration dependent. (A) Superimposed ^1^H-^15^N HSQC spectra of ^15^N-CITED2 CTAD at 100 μM (light blue) or 500 μM (dark blue) protein concentration. Contour levels were maintained at a 1:5 ratio for spectra collected at 100 μM and 500 μM for ease of comparison. Both spectra were collected at 25 °C with identical acquisition parameters in buffer containing 20 mM Tris-HCl pH 6.8, 150 mM NaCl, 2 mM DTT, and 1% D_2_O. (B) Weighted average ^1^H-^15^N chemical shift changes (Δδav) for each residue in the CITED2 CTAD for spectra collected at 100 µM and 500 µM protein concentration. (C) Ratio of peak intensities for CITED2 CTAD residues from spectra collected at different protein concentrations. Intensity ratios were calculated by first dividing the peak intensities from the 500 µM spectrum by a factor of 5 to account for expected protein concentration-dependent increases in peak intensity and then dividing these scaled values by the peak intensities from the 100 µM spectrum. In (B) and (C), data are omitted for resonances that are broadened beyond detection or overlapped in at least one of the spectra. Dashed horizontal lines represent the 10% trimmed mean.

**Figure 5.**
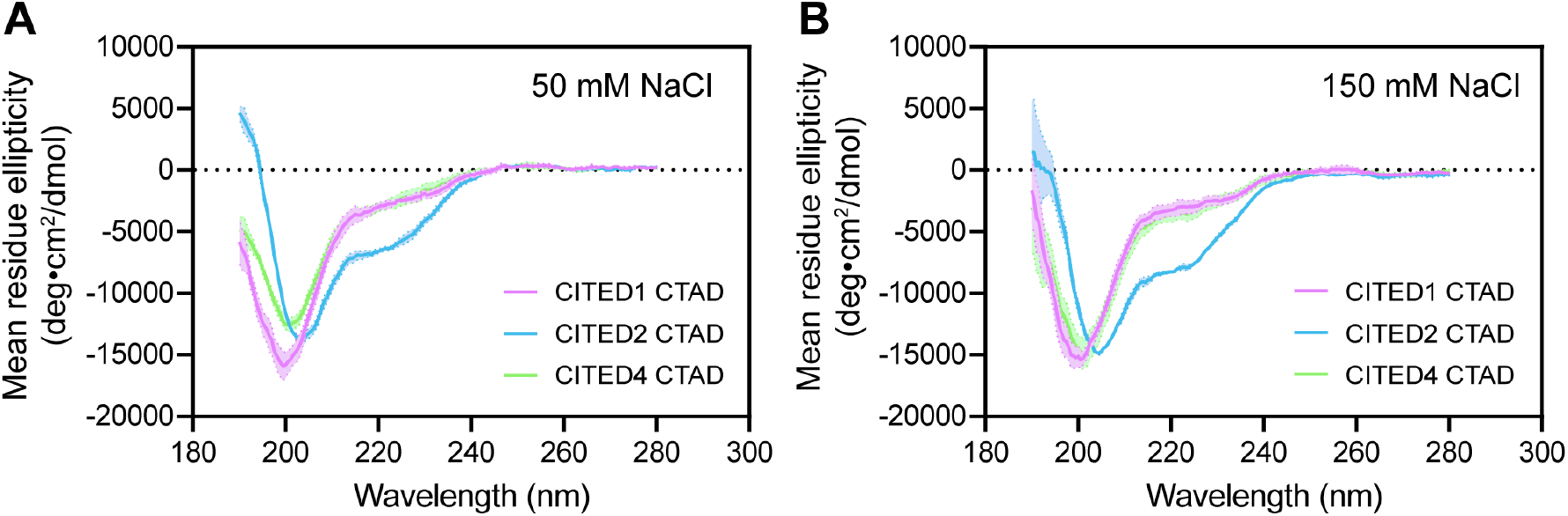
CITED2 exhibits α-helical characteristics by circular dichroism. CD spectra of 100 µM protein samples in buffers containing (A) 5 mM Tris-HCl pH 6.8 and 50 mM NaCl or (B) 5 mM Tris-HCl pH 6.8 and 150 mM NaCl in a 0.01 cm pathlength cuvette at 25 °C. Data for the CITED1 CTAD are shown in light purple, CITED2 CTAD in light blue, and CITED4 CTAD in light green. Shaded regions represent error across three independent measurements.

Similar differences are observed with increasing protein concentration (Figure 4). Again, the most substantial differences in chemical shift and intensity are observed for residues in the *α*A region as well as in the conserved LPEL and ϕC motifs. Notably, we do not observe any changes in peak position in the HSQC spectra of the CITED1 and CITED4 CTADs upon increasing concentration (Supplementary Figure 6). These results highlight that these responses to changes in ionic strength and concentration are specifically encoded in the CITED2 CTAD amino acid sequence.

**Figure 6.**
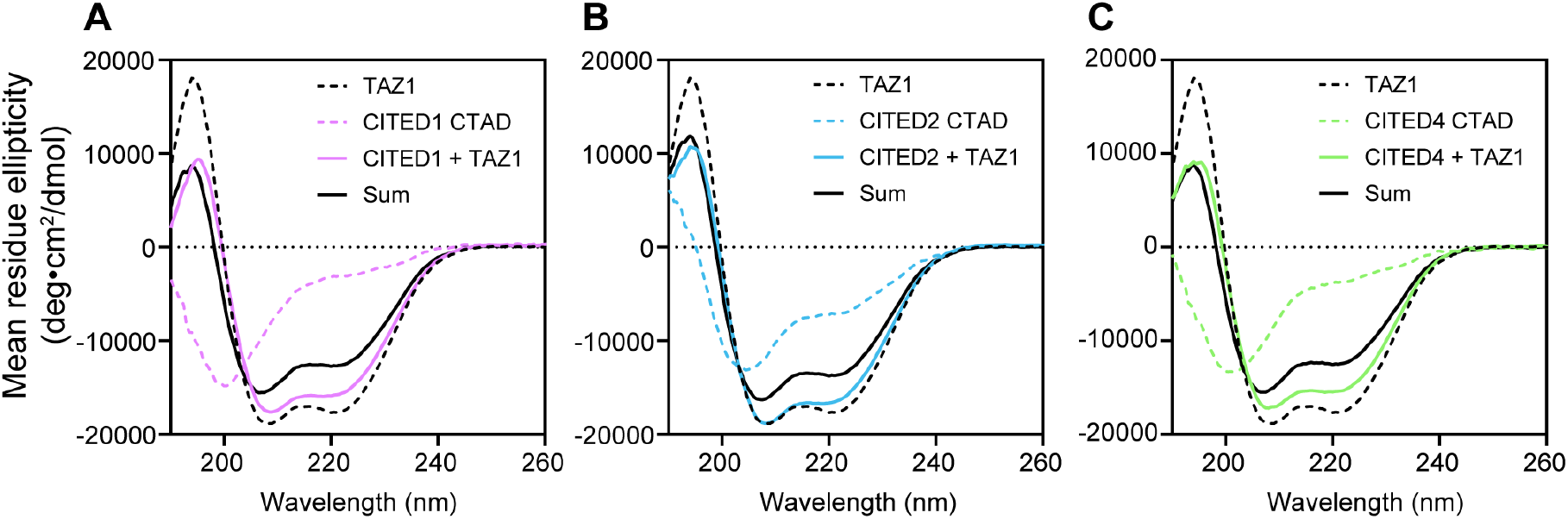
All CITED CTADs undergo coupled folding and binding in complex with TAZ1. Circular dichroism (CD) spectra were collected in buffer containing 5 mM Tris, pH 6.8, 150 mM NaCl for TAZ1 and the CITED CTADs alone and in mixtures for (A) CITED1, (B) CITED2, and (C) CITED4. Individual CD spectra are plotted as described in the inset legends. The sum is calculated as the per residue weighted average of the ellipticities of the individual components (*Materials and Methods*).

Our NMR data indicate that CITED2 CTAD residues in the region that forms an *α* helical structure in complex with TAZ1 are selectively driving the observed spectral changes. To further explore whether differences in secondary structural propensity play a role in driving differences in the CITED2 CTAD conformational ensemble and dynamics, we obtained circular dichroism (CD) spectra for the CITED CTADs. CD spectra of the CITED2 CTAD display marked differences from the spectra for CITED1 and CITED4 (Figure 5). While the CITED1 and CITED4 CTADs largely resemble random coils, the CD spectrum of the CITED2 CTAD suggests that the protein is partially helical, with characteristic minima for *α* helix formation at 208 and 222 nm. CD spectra also confirm that helix formation in the CITED2 CTAD is dependent on the ionic strength of the buffer, with increased helicity observed with increasing NaCl concentration. Thus, there is excellent agreement between the NMR and CD data highlighting that the CITED2 CTAD alone forms a partially populated helical structure in solution.

Prior work has identified that the *α*A helix of CITED2 is essential for high affinity binding to TAZ1^18–20^. Our NMR and CD data for the CITED2 CTAD in isolation highlight that this helix is partially populated in solution, whereas the CITED1 and CITED4 CTADs remain largely disordered. The CITED2 CTAD binds to TAZ1 through a coupled folding and binding mechanism which significantly stabilizes the *α*A helix^16,17^. Given the observed differences in the conformational ensembles of the CITED1 and CITED4 CTADs, we wondered whether these peptides would also interact with TAZ1 in a similar manner. We measured CD spectra for the CITED CTADs and TAZ1 alone and in complex. These data reveal that while the CITED CTADs have distinct conformational ensembles in their unbound states, all of the CTADs undergo coupled folding and binding to adopt partially helical structures in complex with TAZ1 (Figure 6).

In these experiments, the calculated sum of the CD spectra for the free components assumes that there is not a change in secondary structure upon binding. Instead, we observe that the CD spectra of the complexes do not agree with this calculated sum and show decreased ellipticity at 208 nm and 222 nm indicative of a gain in helical structure upon complex formation. Similar behavior was also observed by NMR (Supplementary Figure 7). Mixing the ^15^N-labeled CITED CTADs with unlabeled TAZ1 results in observable peaks over a much wider range of ^1^H chemical shifts that is more consistent with what is typically observed for folded proteins.

**Figure 7.**
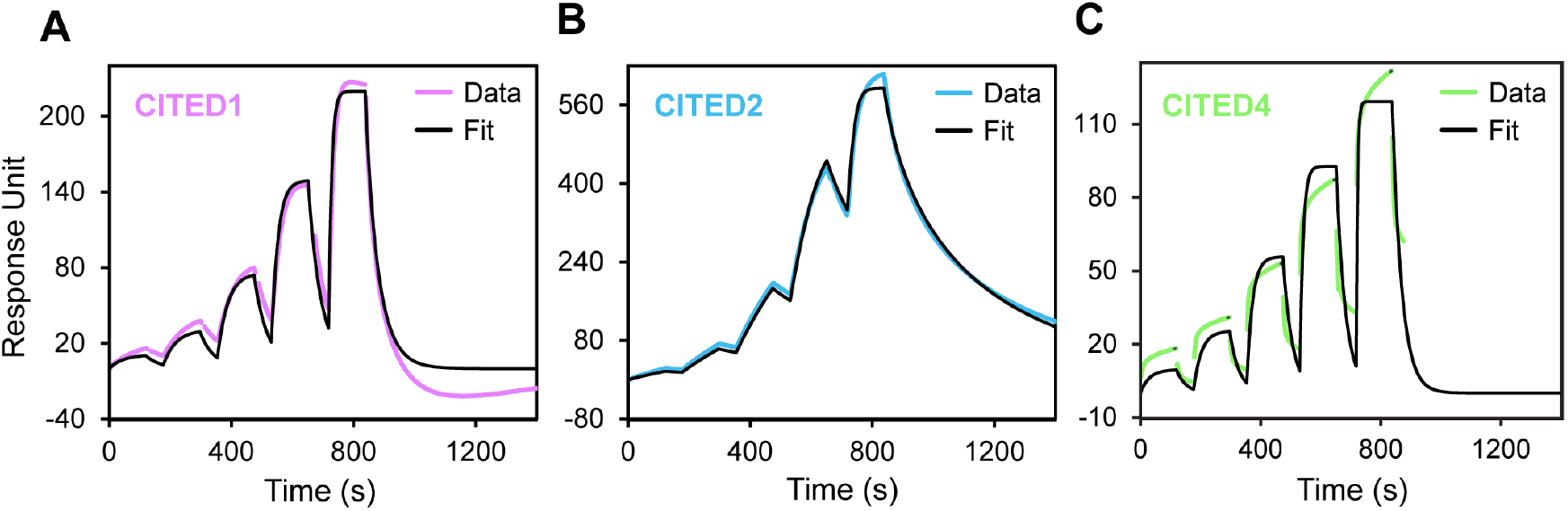
The CITED CTADs bind to TAZ1 with distinct thermodynamics and kinetics. Representative SPR sensorgrams are shown for the (A) CITED1, (B) CITED2, and (C) CITED4 CTADs. Biotinylated TAZ1 was immobilized on the sensor chip, and the CITED1 and CITED2 CTADs were injected at 1.2 nM, 3.7 nM, 11.1 nM, 33.3 nM, and 100 nM concentrations. The CITED4 CTAD was injected at 12 nM, 37 nM, 111 nM, 333 nM, and 1000 nM concentrations. The data were fit to a 1:1 binding model (black line). For the CITED4 CTAD, data after 900s was removed to allow for data fitting.

Together, these results highlight that the CITED CTADs bind to TAZ1 by a common mechanism, which raises a broader question about why their sequences are distinct. Thus, we asked whether variations in sequence impact the binding affinity of the CITED CTADs for TAZ1. We used surface plasmon resonance (SPR) to characterize the thermodynamics and kinetics for the CITED CTAD:TAZ1 interactions. For these experiments, biotinylated TAZ1 (see *Materials and Methods*) was immobilized on a streptavidin sensor chip and increasing concentrations of the CITED CTADs were added sequentially in a single-cycle kinetics experiment. Using this approach, we were able to obtain high-quality binding data that allowed us to extract kinetic and thermodynamic parameters to describe the CITED CTAD:TAZ1 interactions (Figure 7). While all of the CITED CTADs bind to TAZ1 in the nanomolar regime, there are appreciable differences in the affinities of the individual CITED CTADs for TAZ1 that are largely driven through changes in the dissociation rate constant (Table 1 and Figure 7). The *K*_d_ for the CITED2 CTAD:TAZ1 interaction by this method (7 ± 2 nM) agrees well with previously published values determined by other methods^17–19^. The SPR measurements reveal that the CITED1 and CITED4 CTADs have weaker binding affinities for TAZ1 (Table 1). Interestingly, there is a trend in the *K*_d_ values that reflects the number of amino acid substitutions in the CITED1 and CITED4 CTAD sequences relative to CITED2. The sequence of the CITED1 CTAD is more similar to the CITED2 CTAD and binds TAZ1 with slightly weaker affinity, whereas larger differences in affinity are observed for the CITED4 CTAD which has more pronounced differences in its sequence. These differences arise from variations in the kinetic parameters. While the CITED1 and CITED2 CTADs share similar association rate constants, there is an order of magnitude decrease in this parameter for the CITED4 CTAD. The dissociation rate constants reveal that the CITED2 CTAD dissociates from TAZ1 at a slower rate than the CITED1 and CITED4 CTADs. These data highlight that there are features of the CITED CTAD amino acid sequences that are fine-tuning their interactions with their shared binding partner TAZ1 which could be important for distinguishing their functional roles.

**Table 1:** Kinetic and thermodynamic parameters for CITED CTAD:TAZ1 interactions from SPR. Reported values represent the average and standard deviation of at least 3 independent measurements.

| Construct | $k_a$ ($M^{-1}s^{-1}$) | $k_d$ ($s^{-1}$) | $K_D$ (nM) |
| --- | --- | --- | --- |
| CITED1 CTAD | $2.2 \pm 0.9 \times 10^6$ | $4.5 \pm 1.3 \times 10^{-2}$ | $23 \pm 6$ |
| CITED2 CTAD | $2.4 \pm 0.8 \times 10^6$ | $1.5 \pm 0.3 \times 10^{-2}$ | $7 \pm 2$ |
| CITED4 CTAD | $2.2 \pm 0.3 \times 10^5$ | $5.1 \pm 1.6 \times 10^{-2}$ | $220 \pm 40$ |

## Discussion

Recent advances in predicting and modeling conformational ensembles of intrinsically disordered regions have provided much needed insights into sequence-ensemble relationships^23,24^, yet comparative experimental studies are still required to fully understand the relationships between sequence and function in disordered proteins. Our studies of the CITED CTAD sequences in this work add new experimental evidence for the complexities and nuances encoded in the amino acid sequences of disordered activation domains. We find that amino acid differences within the remarkably similar CITED CTADs encode unique molecular features that have measurable impacts on biological function.

Given the degree of similarity of the CITED CTAD sequences in regions that are known to impact CITED2:TAZ1 interactions, we were somewhat surprised to uncover differences in the conformational ensembles of the CITED CTADs in their unbound state. AGADIR predictions^25–28^ hint at potential differences in the CTAD conformational ensembles but do not agree entirely with our experimental results. AGADIR predicts ~7% helicity for the CITED2 *α*A helix (residues 225-235) in isolation, whereas the corresponding region in CITED1 (residues 155-165) is predicted to be ~5% helical. While this is potentially in agreement with our experimental results, AGADIR also predicts that the corresponding region in CITED4 (residues 139-149) is ~7% helical. Our CD experiments reveal that the CITED2 CTAD has more helical content than the other CTADs (Figure 5). The differences between the CTAD sequences but high degree of conservation for a given member of the family suggests that there is some selective pressure to preserve the sequence differences in the CITED CTADs.

The differences in the CITED CTAD sequences may have been preserved across evolution to differentiate interactions with the TAZ1 domain of CBP/p300 or other coactivators. Our data highlight that the different CTAD sequences have different binding affinities for TAZ1, with the CITED2 CTAD sequence seemingly being optimal for high affinity binding (Figure 7 and Table 1). The effects of amino acid substitutions can be interpreted from the solution structure of the CITED2:TAZ1 complex, where many residues on the hydrophobic face of the amphipathic *α*A helix make stabilizing contacts with the TAZ1 *α*1 and *α*4 helices^17^. These residues (I223, V227, M229, V232, and M235) are substituted with amino acids with smaller or more polar sidechains in the CITED1 and/or CITED4 sequences (Figure 1), suggesting that the diminished binding affinities for TAZ1 for these sequences is due to less favorable interactions of the CITED1 and CITED4 CTADs in complex with TAZ1. A comprehensive mutational study of human activation domain sequences identified that the CITED2 CTAD is an exceptionally “strong” activation domain and that mutations in highly conserved regions of the CITED2 CTAD sequence attenuate transcriptional activation^29^. The positions with the largest loss of activation upon mutation are positions that are conservatively substituted in the CITED1 and CITED4 CTADs. For example, substituting conserved aspartate residues towards the N-terminus of the CITED2 CTAD (D221 and D224) with glutamate residues resulted in a decrease in transcriptional activation activity. While D224 is conserved across the human CITED CTADs, the position corresponding to D221 in CITED4 is a glutamate. Another example that highlights these effects is mutation of W247 to phenylalanine, which also decreases activity in the prior study. Again, this position is a conserved tryptophan in both CITED1 and CITED2 but is substituted with a phenylalanine residue in the CITED4 CTAD.

Our NMR data for the CTADs highlight that the CITED2 CTAD is more structurally and/or dynamically restricted than the CITED1 and CITED4 CTADs. The increase in helicity observed with increasing ionic strength (Figure 5) agrees with prior studies of helical peptides^25,30,31^. Residues in the *α*A region of the CITED2 CTAD are most impacted by increasing concentration, suggesting that the concentration-induced broadening that we observe could be due to reversible helix self-association^32^. As the CITED2 *α*A helix is amphipathic, it is reasonable to expect that helix self-association could be energetically favorable to shield the hydrophobic face of the helix from solvent, particularly at the high concentrations used for our experiments. We also cannot rule out the presence of additional helix-stabilizing tertiary interactions from the data presented here, although exciting studies of CITED2 CTAD behavior in cells suggest that CITED2 can adopt more compact states in comparison to other transcriptional activation domains^33^. Additional experiments will be needed to fully ascertain the origins of the complex structural and dynamic behavior unique to the CITED2 CTAD.

The observed residual helicity in the CITED2 CTAD may be important for its binding mechanism and could contribute to its higher binding affinity for TAZ1 in comparison with the other CTADs. Studies of other IDPs have highlighted that disrupting pre-formed structures in the conformational ensemble can significantly alter binding free energies and kinetics as well as biological functions^34–37^. While the data presented here demonstrate a correlation between helicity and high affinity binding, further studies will be required to fully understand these relationships. Importantly, as the *α*A region in the CITED2 CTAD is known to have a critical role in allowing CITED2 to rapidly displace the HIF-1*α* CTAD from TAZ1^18,19^, it will be interesting to investigate whether the CITED1 and CITED4 CTADs have diminished ability to compete with HIF-1*α* CTAD or other transcriptional activation domains for TAZ1 binding. We speculate that substitutions in the CITED1 and CITED4 CTADs alter the valency of interactions with TAZ1 that would in turn result in less effective competition for TAZ1 binding^7,18,19,38^.

In summary, the atomic resolution data presented here coupled with mechanistic biophysical characterization of the CITED CTADs and their interactions with TAZ1 provide a new framework for understanding how subtle sequence substitutions can have differing effects on IDP structures, dynamics, and functions. Our studies of the human CITED CTADs and their conformational ensembles and interactions reveal exciting insights into sequence variation across protein families that emphasize the importance of such studies for fully capturing the functional intricacies of IDPs.

## Materials and Methods

### Constructs

The DNA sequences encoding for the human CITED1 (residues 146-193) and CITED4 (residues 130-184) C-terminal transactivation domains were cloned into a modified bicistronic pET22b vector designed for co-expression of His_6_-tagged GB1 fusion proteins with the TAZ1 domain of mouse CBP (residues 340-439)^39^. DNA for CITED1 was synthesized by Twist Biosciences. CITED4 cDNA was purchased from OriGene Technologies. The vector for co-expression of the His_6_-GB1-CITED2 CTAD (residues 216-270) and the TAZ1 domain of mouse CBP (residues 340-439) was generated using site-directed mutagenesis to introduce a codon for the native C-terminal residue Cys270 into a previously generated plasmid encoding human CITED2 residues 216-269 (Addgene #173761)^18^. In each of these vectors, the His_6_-GB1 fusion sequence is followed by a thrombin cleavage site. For CITED2, there are five non-native residues (GSHMS) at the N-terminus of the CTAD sequence after cleavage. The CITED1 and CITED4 CTADs have four non-native residues (GSHM) after cleavage.

A pET21d vector for expression of the TAZ1 domain of mouse CBP (residues 340-439) was purchased from Addgene (#173760).^40^ A construct of TAZ1 with N-terminal His_6_ and Avi-tag sequences (GLNDIFEAQKIEWHE) (Avi-TAZ1) in a pET21(+) vector was synthesized by Twist Biosciences.

### Protein Expression and Purification

The CITED CTADs were expressed in E. coli BL21 (DE3) as His_6_-tagged GB1 fusion proteins in a co-expression vector with TAZ1 (residues 340–439 of mouse CBP). Initial steps of the purification protocol (lysis, nickel affinity chromatography, and thrombin cleavage) were carried out as previously described.^18,41^ Following overnight thrombin cleavage on Ni-NTA resin at 4 °C, the cleaved CITED CTAD peptides were washed off the resin using buffer containing 20 mM Tris pH 8 and subsequently loaded onto a HiTrap™ Q HP anion exchange chromatography column (Cytiva) equilibrated in the same buffer. The CITED CTAD peptides were eluted from the column over a gradient to 1M NaCl. Pure CITED CTAD peptides were flash frozen and stored at −80°C.

TAZ1 was expressed and purified under native conditions as described previously.^18^ To obtain biotinylated TAZ1 for surface plasmon resonance experiments, soluble Avi-TAZ1 was purified by nickel affinity chromatography using Ni-NTA resin. Biotinylation of Avi-TAZ1 was achieved using the BirA biotin-protein ligase standard reaction kit from Avidity. Biotinylated TAZ1 was further purified using size-exclusion chromatography on a Superdex 75 column (Cytiva). The extent of protein biotinylation was determined using a streptavidin gel-shift assay as previously described.^42^

All proteins were expressed in M9 minimal media supplemented with 100 µM ZnSO_4_ at induction. Uniform isotopic labeling for NMR samples was achieved by expression in M9 minimal media supplemented with ^13^C glucose (2.5 g/L) and/or ^15^N ammonium sulfate (1.0 g/L) as the sole carbon and nitrogen sources, respectively.

### Concentration Determination

Protein concentrations were determined by absorbance at 280 nm. For the CITED4 CTAD, which does not contain any tryptophan or tyrosine residues, concentrations were determined by absorbance at 205 nm^43^. To ensure agreement between concentrations determined from absorbance measurements at 280 nm and 205 nm, we measured the absorbance at both wavelengths for proteins containing tryptophan and tyrosine residues and found excellent agreement between the calculated protein concentrations, with less than 5% variation between concentrations determined by absorbance at 280 nm and 205 nm.

### NMR Spectroscopy

NMR samples were prepared in buffer containing 20 mM Tris-HCl pH 6.8, 50 mM or 150 mM NaCl, 2 mM DTT, and 1% D_2_O. All spectra were collected at 25 °C on Bruker Avance III HD 700 and 850 MHz spectrometers equipped with TCI cryoprobes. NMR data were processed using NMRPipe^44^ and analyzed using NMRFAM-SPARKY^45^ on the NMRbox platform^46^.

Backbone chemical shift assignments for the CITED2 CTAD were obtained using standard triple-resonance experiments (HNCO, HNCACB, and HN(CO)CACB) as implemented in the Bruker pulse program library (TopSpin version 3.6.5). Assignment data were collected for a sample of 100 µM ^13^C,^15^N-labeled CITED2 CTAD in 20 mM Tris-HCl pH 6.8, 50 mM NaCl, 2 mM DTT, 1% D_2_O. The indirect dimensions of the HNCACB and HN(CO)CACB spectra were acquired using non-uniform sampling (10%). Non-uniformly sampled spectra were reconstructed using SMILE^47^.

Assignments were transferred to ^1^H-^15^N HSQC spectra collected at different protein and salt concentrations by comparison. For comparisons between ^1^H-^15^N HSQC spectra, weighted average proton and nitrogen chemical shift changes are calculated as Δδ_av_ = ((Δδ_H_)^2^ + (Δδ_N_/5)^2^)^1/2^.

### Surface Plasmon Resonance (SPR)

All SPR experiments were conducted at 25 °C on a Biacore 8K instrument equilibrated in 20 mM Tris-HCl pH 6.8, 150 mM NaCl, 0.05% Tween 20, and 2 mM DTT (running buffer). All proteins were dialyzed at 4 °C overnight into running buffer prior to use. Biotinylated TAZ1 was diluted to 100 nM in running buffer and immobilized onto a Series S streptavidin (SA) Sensor Chip (Cytiva) with a contact time of 60 s at a flow rate of 10 µL/min to yield a Rmax of ~600–800 RU. Following immobilization, affinity measurements were made with five 1:3 serial dilutions of each CITED CTAD using a single-cycle kinetics method. Data for the CITED1 and CITED2 CTADs were obtained using analyte concentrations from 1.2 nM to 100 nM, and data for the CITED4 CTAD was obtained using analyte concentrations from 12 nM to 1 μM. Data were collected using both flow cells and five concentrations per cycle with the following experimental parameters: contact time, 120 s; dissociation time, 600 s; flow rate, 30 μL/min. All data were analyzed with the Biacore Insight Evaluation software (version 3.0.12) using the general single-cycle kinetics method and were fit to the 1:1 binding kinetics model. All experiments were repeated at least three times, using a new sensor chip and fresh protein preparations for each replicate.

### Circular dichroism (CD)

CD spectra were collected at 25 °C using a JASCO J-815 CD spectrometer and a 0.1 mm pathlength cuvette. All samples were prepared in buffer containing 5 mM Tris-HCl pH 6.8 and 50 mM or 150 mM NaCl at a protein concentration of 100 µM. Coupled folding and binding experiments were carried out with mixtures of the CITED CTADs and TAZ1 at a 1:1 molar ratio. CD spectra were collected in continuous scanning mode from 190 – 280 nm with 50 nm/min data pitch for three accumulations. All CD spectra were baseline corrected using spectra collected for samples of the corresponding buffer.

Mean residue ellipticity (MRE) was calculated from the raw ellipticity θ using the equation:

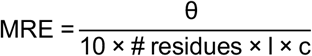

where MRE is in units of deg cm^2^/dmol, θ is in units of mdeg, l is the pathlength of the cuvette in cm, and c is the protein concentration in M.

For coupled folding and binding experiments of protein mixtures, the sum of the CD spectra of the individual components can be calculated as the per residue weighted average of the ellipticities of the individual components:

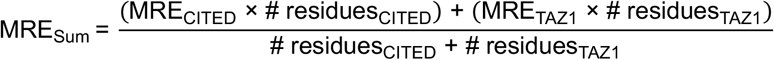

This equation assumes that there is no structural change upon binding for either of the components.

### Sequence Alignments

Amino acid sequences for CITED1, CITED2, and CITED4 CTADs were identified by a BLAST search of the full-length proteins using the Uniprot Database (accessed on December 17, 2025). Sequences were aligned using Clustal Omega^48^ using default settings and visualized using Jalview^49^ (jalview.org). Species names are abbreviated using the format *Homo sapiens* to *H. sapiens*. Accession numbers for all sequences are listed in Supplementary Table 1.

## Acknowledgments

We gratefully acknowledge all members of the Berlow Lab for their critical feedback and helpful discussions about this work. We also thank Dr. Brian Kuhlman and James McGuire Metts for sharing reagents and protocols and providing instrument access, training, and assistance with CD experiments; Dr. Ashutosh Tripathy and Dr. Christopher Travis from the Macromolecular Interactions Core for assistance with SPR experiments; and Dr. Frank Appling, Dr. Jane Dyson, and Dr. Peter Wright for sharing preliminary assignments for a similar construct of the CITED2 CTAD. The UNC Biomolecular NMR Laboratory and the UNC Macromolecular Interactions Facility receive funding from the National Cancer Institute of the NIH under award number P30CA016086. This study made use of NMRbox: National Center for Biomolecular NMR Data Processing and Analysis, which is supported by NIH grant P41GM111135 (NIGMS). This work was generously supported by funds from the Department of Biochemistry and Biophysics and the Dean’s Office at the UNC School of Medicine as well as the Lineberger Comprehensive Cancer Center.

## Data Availability

NMR backbone chemical shift assignments have been deposited in the BMRB under accession number 53509. All plasmids generated for this study are available on Addgene. All other data and materials are available upon request.

