## Supplementary Material for "Sequence-encoded differences in the conformational ensembles of CITED transcriptional activation domains impact coactivator binding"

```
Human H.sapiens/146-193              AIIDSDPVDEEVLMSLVVELGLDRANELPELWLGQNEFDFTADFPSSC
Mouse M.musculus/156-203             ALIDSDPVDEEVLMSLVVELGLDRANELPELWLGQNEFDFTADFPSGC
Bovine B.taurus/148-195              AIIDSDPVDEEVLMSLVVELGLDRANELPELWLGQNEFDFTADFPSGS
Horse E.caballus/148-195             AIIDSDPVDEEVLMSLVVELGLDRANELPELWLGQNEFDFTADFPSGC
Dog C.lupus/174-221                  AIIDSDPVDEEVLMSLVVELGLDRANELPELWLGQNEFDFTADFPSGC
Elephant L.africana/126-173          AIIDSDPVDEEVLMSLVVELGLDQASDLPELWLGQNEFDFTADFATGC
Platypus O.anatinus/140-187          GIIDSDPVDEEVLMSLVVELGLDRANELPELWLGQNEFDFAADFPSGC
Coelacanth L.chalumnae/118-165       GIIDSDPVDEDVLMSLVMELGLDRVDELPELWLGQNEFDFISDLSSGC
```

**Supplementary Figure 1. Sequence alignment of CITED1 CTAD sequences from representative vertebrate and invertebrate species.** Identical residues among sequences are bolded and highlighted in a darker shade of blue. Biochemically conserved residues are highlighted in a lighter shade of blue.

```
Human H.sapiens/216-270           NVIDTDFIDEEVLMSLVIEMGLDRIKELPELWLGQNEFDFMTDFVCKQQPSR--VSC
Mouse M.musculus/215-269          NVIDTDFIDEEVLMSLVIEMGLDRIKELPELWLGQNEFDFMTDFVCKQQPSR--VSC
Bovine B.taurus/219-273           NVIDTDFIDEEVLMSLVIEMGLDRIKELPELWLGQNEFDFMTDFVCKQQPSR--VSC
Horse E.caballus/217-271          NVIDTDFIDEEVLMSLVIEMGLDRIKELPELWLGQNEFDFMTDFVCKQQPSR--VSC
Dog C.lupus/226-280               NVIDTDFIDEEVLMSLVIEMGLDRIKELPELWLGQNEFDFMTDFVCKQQPSR--VSC
Bat D.rotundus/218-272            NVIDTDFIDEEVLMSLVIEMGLDRIKELPELWLGQNEFDFMTDFVCKQQPSR--VSC
Pangolin M.pentadactyla/214-268   NVIDTDFIDEEVLMSLVIEMGLDRIKELPELWLGQNEFDFMTDFVCKQQPSR--VSC
Elephant L.africana/217-271       NVIDTDFIDEEVLMSLVIEMGLDRIKELPELWLGQNEFDFMTDFVCKQQPSR--VSC
Platypus O.anatinus/195-249       NVIDTDFIDEEVLMSLVVEMGLDRIKELPELWLGQNEFDFMTDFVCKQQPSR--VSC
Chicken G.gallus/182-236          NVIDTDFIDEEVLMSLVIEMGLDRIKELPELWLGQNEFDFMTDFVCKQQPSR--VSC
Frog X.tropicalis/171-225         SVIDTDFIDEEVLMSLVIEMGLDRIKELPELWLGQNEFDFMTDFVCKQQPNR--VSC
Lungfish P.annectens/198-252      NVIDTDFIDEEVLMSLVVEMGLDRIKELPELWLGQNEFDFMTEFVCKQQPSR--VSC
Shark C.carcharias/207-261        NVIDTDFIDEEVLMSLVIELGLDRIKELPELWLGQNEFDFMTDFVCKQQPSR--VSC
Ray M.hypostoma/185-239           NVIDTDFIDEEVLMSLVIELGLDRIKELPELWLGQNEFDFMTEFVCKQQPSR--VSC
Coelacanth L.chalumnae/175-225    NVIDTDFIDEEVLMSLVIEMGLDRIKELPELWLGQNEFDFIHDGLCLQTAA------
Zebrafish D.rerio/168-222         NVIDTDFIDEEVLMSLVIEMGLDRIKELPELWLGQNEFDFMTDFVCKQQPSR--VSC
Carp C.carassius/172-226          NVIDTDFIDEEVLMSLVIEMGLDRIKELPELWLGQNEFDFMTDFVCKQQPSR--VSC
Trout O.mykiss/180-234            NVIDTDFIDEEVLMSLVIEMGLDRIKELPELWLGQNEFDFMTDFVCKQQPSR--VSC
Salmon S.salar/180-234            NVIDTDFIDEEVLMSLVIEMGLDRIKELPELWLGQNEFDFMTDFVCKQQPSR--VSC
Lamprey P.marinus/220-266         TMPNADALDEELLMSLVLELGLDRVQELPELWLGQDEVDFLSGVYAH----------
Lancelet B.floridae/277-330       GLSDLDMIDEDILARLIIELGLDRMRELPELWLSHNELEMD---ICRQLPPQRAVRC
Lancelet B.lanceolatum/274-327    GLSDLDLIDEDILARLIIELGLDRMRELPELWLSHNELEMD---ICRQVPPQRAVRC
Lancelet B.belcheri/273-326       GLSDLDMIDEDILARLIIELGLDRMRELPELWLSHNELEMD---ICRQVPPQRAVRC
```

**Supplementary Figure 2. Sequence alignment of CITED2 CTAD sequences from representative vertebrate and invertebrate species.** Identical residues among sequences are bolded and highlighted in a darker shade of blue. Biochemically conserved residues are highlighted in a lighter shade of blue.

| | |
|---|---|
| ***Human H.sapiens/130-184*** | GGMDAELIDEEALTSLELELGLHRVRELPELFLGQSEFDCFSDLGSAPPAGSVSC |
| ***Mouse M.musculus/128-182*** | GCMDTELIDEEALTSLELELGLHRVRELPELFLGQSEFDCFSDLGSAPAAGSVSC |
| ***Bovine B.taurus/131-185*** | GGMDAELIDEEALTSLELELGLHRVRDLPELFLGQSEFDCFSDLGSAPPAGSVSC |
| ***Horse E.caballus/177-231*** | GGMDAELIDEEALTSLELELGLHRVRELPELFLGQSEFDCFSDLGSAPPAGSVSC |
| ***Dog C.lupus/127-181*** | GCMDAELIDEEALTSLELELGLHRVRELPELFLGQSEFDCCSDLGSAPPAGSVSC |
| ***Elephant L.africana/86-140*** | VGMDAELIDEEALTSLELELGLHRVRELPELFLGQSEFDCFSDLGSAPPAGSVSC |
| ***Koala P.cinereus/141-195*** | SCVDAELIDEETLTSLEQELGLDRVQELPELFLGQNEFDCLWDFGGKQQAGAVSC |
| ***Platypus O.anatinus/149-203*** | NLMDTELVDEEVLTTLALELGLDREQELPELFLGQNEFDLVTDLAGKQQAGAVRC |

**Supplementary Figure 3. Sequence alignment of CITED4 CTAD sequences from representative vertebrate and invertebrate species.** Identical residues among sequences are bolded and highlighted in a darker shade of blue. Biochemically conserved residues are highlighted in a lighter shade of blue.

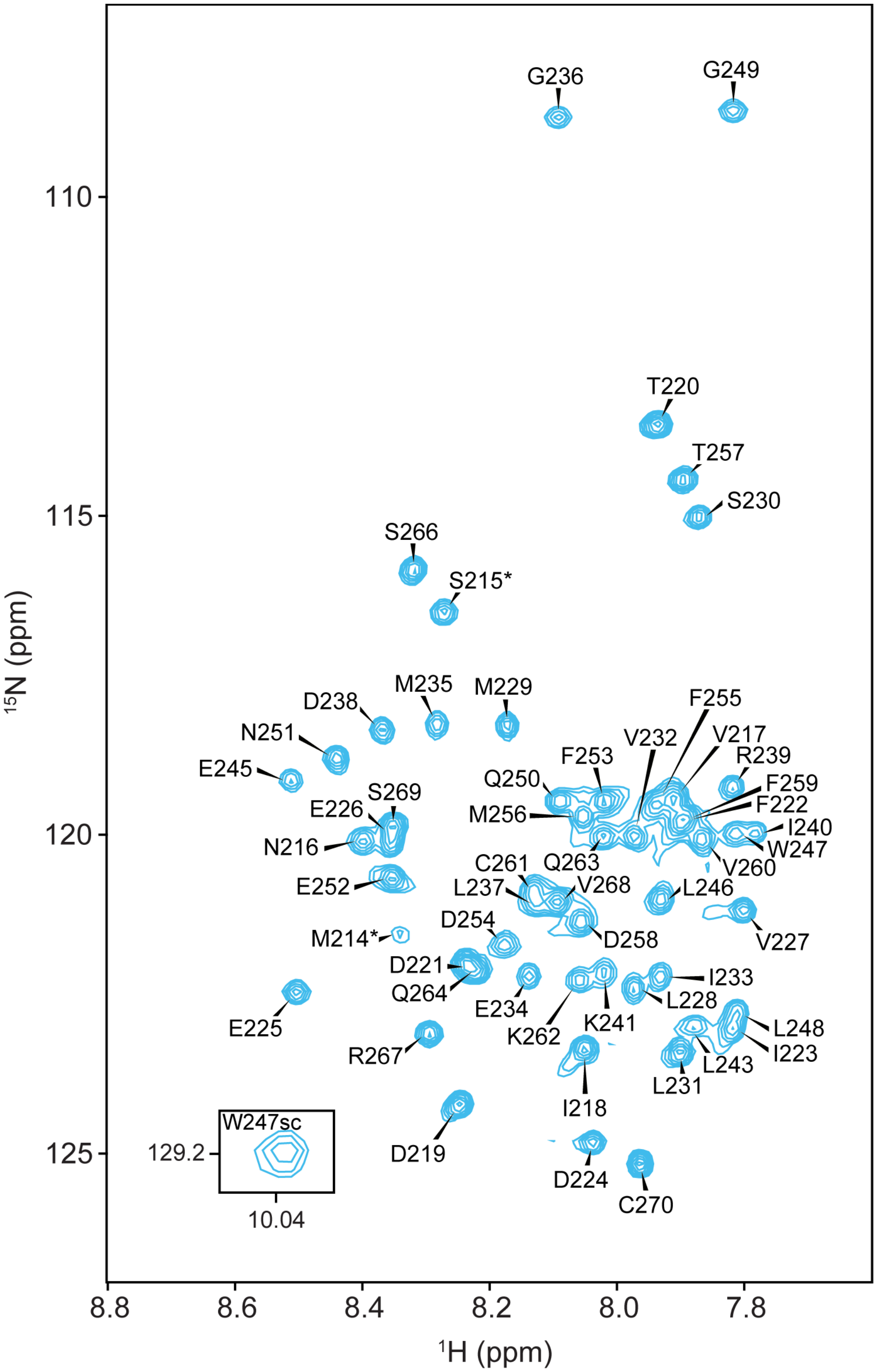


**Supplementary Figure 4. Assigned $^{1}H$-$^{15}N$ HSQC spectrum of the CITED2 CTAD.** Peaks with validated resonance assignments (BMRB entry 53509) are labeled on a representative HSQC spectrum of 100 μM $^{15}N$-labeled CITED2 CTAD in 20 mM Tris-HCl pH 6.8, 50 mM NaCl, 2 mM DTT, 1% $D_2O$. The tryptophan indole sidechain peak is shown as an inset. Asterisks indicate resonances corresponding to non-native residues as described in the *Materials and Methods*.

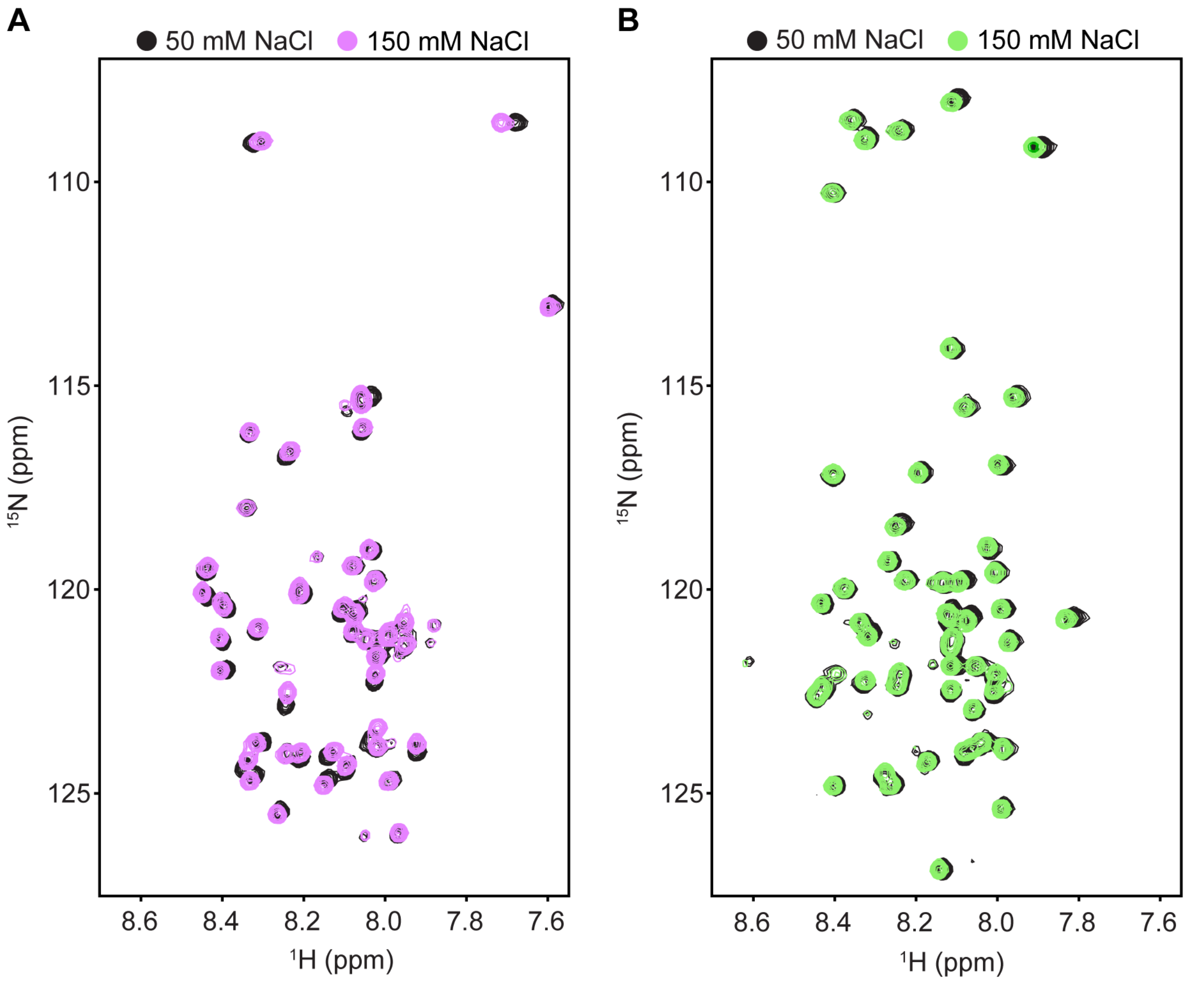


**Supplementary Figure 5. Effect of ionic strength on the $^{1}$H-$^{15}$N HSQC spectra of CITED1 and CITED4.** (A) Superimposed $^{1}$H-$^{15}$N HSQC spectra of 100 μM $^{15}$N-CITED1 CTAD in buffer containing 50 mM NaCl (black) or 150 mM NaCl (light purple) and 20 mM Tris-HCl pH 6.8, 2 mM DTT, and 1% $D_2O$. (B) Superimposed $^{1}$H-$^{15}$N HSQC spectra of 100 μM $^{15}$N-CITED4 CTAD in buffer containing 50 mM NaCl (black) or 150 mM NaCl (light green) and 20 mM Tris-HCl pH 6.8, 2 mM DTT, and 1% $D_2O$. All spectra were collected at 25 °C.

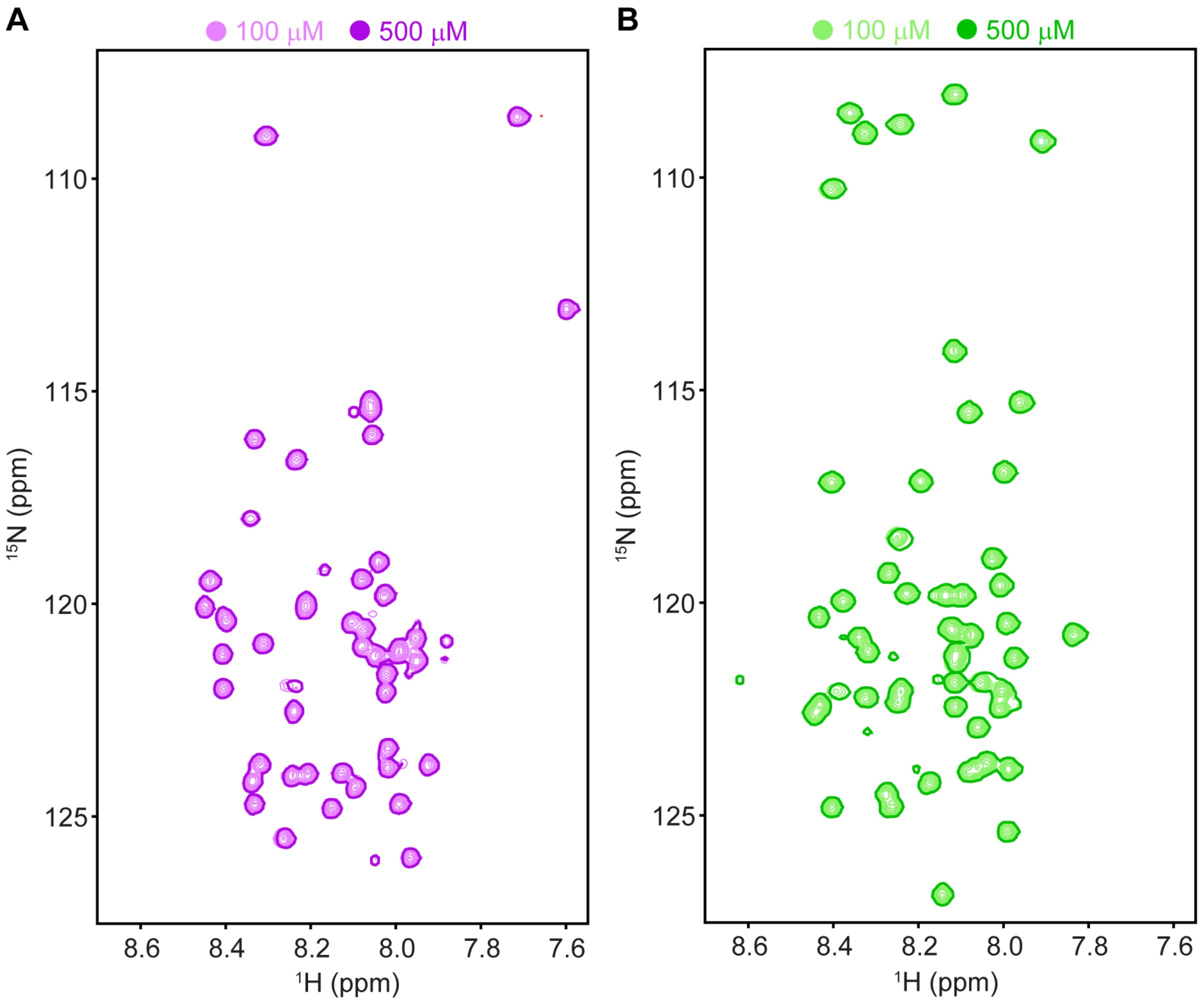


**Supplementary Figure 6. Effect of protein concentration on the $^{1}$H-$^{15}$N HSQC spectra of CITED1 and CITED4.** (A) Superimposed $^{1}$H-$^{15}$N HSQC spectra of $^{15}$N-CITED1 CTAD at 500 μM (dark purple, fewer contours) or 100 μM (light purple) protein concentration. (B) Superimposed $^{1}$H-$^{15}$N HSQC spectra of $^{15}$N-CITED4 CTAD at 500 μM (dark green, fewer contours) or 100 μM (light green) protein concentration. For (A) and (B), contour levels were maintained at a 1:5 ratio for spectra collected at 100 μM and 500 μM for ease of comparison. Fewer contours are shown for the 500 μM spectra due to the high degree of similarity. Both spectra were collected at 25 °C with identical acquisition parameters in buffer containing 20 mM Tris-HCl pH 6.8, 150 mM NaCl, 2 mM DTT, and 1% $D_2O$.

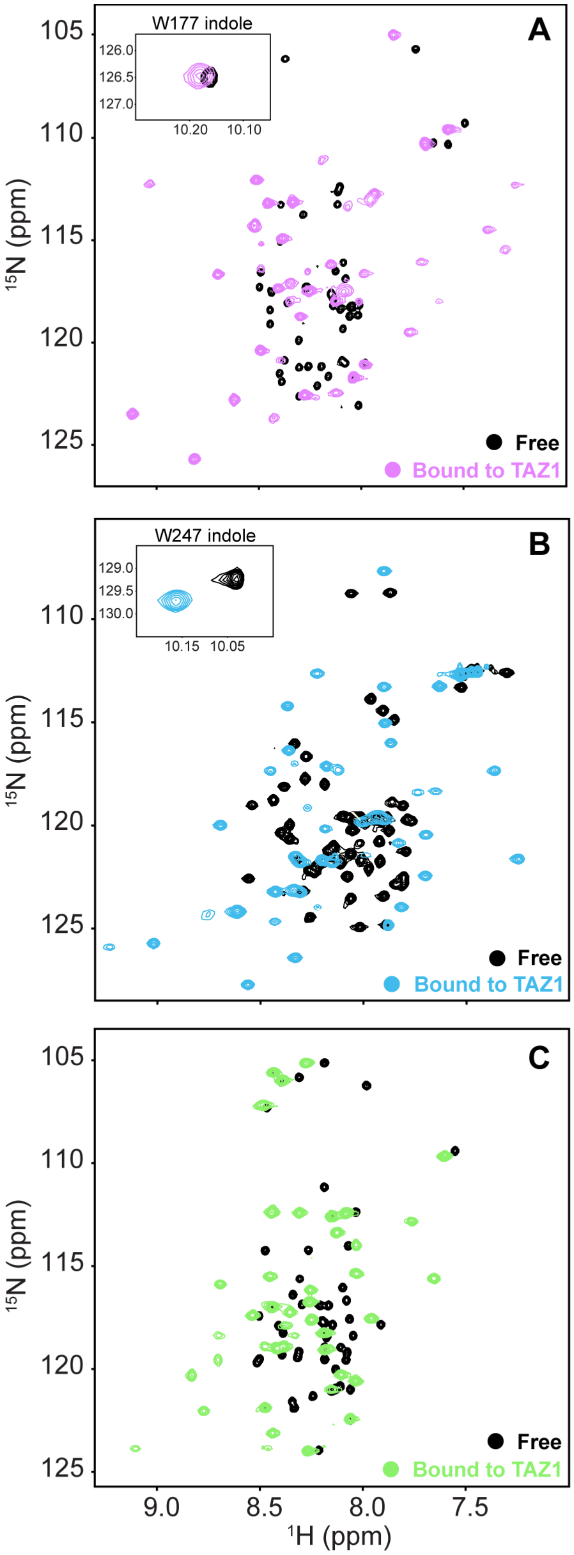


**Supplementary Figure 7.** The CITED CTADs undergo coupled folding upon binding in complex with TAZ1. (A-C) Superimposed $^{1}H$-$^{15}N$ HSQC spectra for the $^{15}N$-CITED CTADs alone (black) or in complex with TAZ1. All spectra were collected at 25 °C in buffer containing 20 mM Tris-HCl pH 6.8, 150 mM NaCl, 2 mM DTT, and 1% $D_2O$.
